# Megabase-Scale Transgene De-Duplication to Generate a Functional Single-Copy Full-Length Humanized DMD Mouse Model

**DOI:** 10.1101/2024.03.25.586713

**Authors:** Yu C. J. Chey, Mark A. Corbett, Jayshen Arudkumar, Sandra G. Piltz, Paul Q. Thomas, Fatwa Adikusuma

## Abstract

The development of sequence-specific precision treatments like CRISPR gene editing therapies for Duchenne Muscular Dystrophy (DMD) requires sequence humanised animal models to enable the direct clinical translation of tested strategies. The current available integrated transgenic mouse model containing the full-length human *DMD* gene, Tg(DMD)72Thoen/J (hDMDTg), has been found to have two copies of the transgene per locus in a tail-to-tail orientation, which does not accurately simulate the true copy number of the *DMD* gene. This duplication also complicates analysis when testing CRISPR therapy editing outcomes, as large genetic alterations and rearrangements can occur between the cut sites on the two transgenes. To address this, we performed long read nanopore sequencing on hDMDTg mice to better understand the structure of the duplicated transgenes. Following that, we performed a megabase-scale deletion of one of the transgenes by CRISPR zygotic microinjection to generate a single-copy, full-length, humanised DMD transgenic mouse model (hDMDTgSc). Functional, molecular, and histological characterisation shows that the single remaining human transgene retains its function and rescues the dystrophic phenotype caused by endogenous murine *Dmd* knockout. Our unique hDMDTgSc mouse model can potentially be used for the further generation of DMD disease models that would be better suited for the pre-clinical assessment and development of sequence specific CRISPR therapies.

## Introduction

Duchenne Muscular Dystrophy (DMD) is an X-linked monogenic disorder characterised by progressive wasting of skeletal and heart muscles [1]. This condition has an incidence of around 1 in 5,000 newborn boys and is caused by loss-of-function mutations in the X-linked *DMD* gene, resulting in the absence of dystrophin protein expression [2, 3]. *DMD* is the largest human gene, encompassing 79 exons spanning 2.4 Mb. It encodes various short and long dystrophin isoforms that are differentially expressed in muscle and the nervous system [4]. The predominant muscle isoform, Dp427m, plays a crucial role in connecting intracellular actin filaments with extracellular laminin via the dystrophin associated glycoprotein complex, which is essential for maintaining and stabilising the muscle cell membrane (sarcolemma) during contractions. Without dystrophin, the sarcolemma is prone to contraction-induced mechanical stress and damage, leading to the progressive replacement of muscle with fat and fibrous tissue [5].

Most DMD-causing mutations are large exonic deletions clustering around deletion hotspots that disrupt the reading frame [6]. Mutations that otherwise retain the *DMD* reading frame are associated with the less common and symptomatically milder Becker Muscular Dystrophy (BMD), where internally truncated but partially functional dystrophin protein is produced.

Pre-clinical testing of sequence-specific therapies for DMD, such as antisense oligonucleotide (ASO) and CRISPR gene editing, require humanised animal models due to variations in the *DMD* gene sequence across different species [7]. Strategies designed to target endogenous animal sequences may not necessarily retain their efficiencies or specificities when converted to their corresponding human-targeting sequence. Thus, humanised animal models that incorporate the entire human *DMD* genomic sequence are invaluable for assessing patient-relevant therapeutic candidates that target intronic sequences or modulate splicing.

The first humanised DMD mouse model with stable germline transmission, Tg(DMD)72Thoen/J (hDMDTg), was created by integrating a yeast artificial chromosome (YAC) containing the full-length human *DMD* (*hDMD*) transgene into chromosome 5 of mouse embryonic stem (ES) cells [8]. When crossed with the *mdx* strain lacking murine dystrophin expression, the resulting hDMDTg/*mdx* mice exhibited functional compensation for the absence of mouse dystrophin through human dystrophin expression. Because it possesses the complete human *DMD*, including proximal regulatory sequences (promoter), exons and introns, the hDMDTg/*mdx* strain serves as a platform for the further creation of humanised DMD disease models by the introduction of mutations into the *hDMD* transgene that mimic various DMD patient mutations. Examples of these mouse disease models include the exon 52 deletion hDMDTgEx52Δ/*mdx* models [9, 10] and the exon 45 deletion hDMDTgEx45Δ/*mdx* and hDMDTgEx45Δ/*mdxD2* models [11]. A second mouse model with the full length human *DMD* gene carried on a non-integrated human artificial chromosome (DYS-HAC) has also been generated, but mice with DYS-HACs containing DMD mutations has not yet been developed [12, 13].

During the detailed characterisation of their TALEN-generated hDMDTgEx52Δ/*mdx* mouse model, Yavas *et al.* (2020) identified a tail-to-tail duplication of the *hDMD* transgene present in both the hDMDTgEx52Δ/*mdx* and hDMDTg/*mdx* mice by interphase FISH and digital droplet PCR, indicating that a duplication event likely occurred during the initial generation of the hDMDTg model [14]. The two-copy number of this transgene does not accurately reflect the single-copy nature of the *DMD* gene in males and also makes the generation of mouse models harbouring patient mutations much more complicated.

In this study, we carried out a detailed genetic analysis of hDMDTg mice using short- and long-read whole genome sequencing to resolve the structure of the duplicated hDMD transgenes. We then de-duplicated the *hDMD* transgene in hDMDTg mice through the CRISPR microinjection of zygotes to generate a single-copy hDMD mouse strain, hDMDTgSc. Additionally, we provide characterisation of these hDMDTgSc mice and show that the single remaining transgene copy is functional and rescues the phenotype associated with the endogenous *Dmd* knock-out mouse.

## Results

### Sequence analysis of the hDMDTg model with duplicated hDMD transgenes

To identify the 5’ and 3’ boundaries of the *DMD* locus of the hDMDTg transgene, short-read genome sequencing was performed on genomic DNA extracted from the liver of homozygous hDMDTg mice. The sequencing data was mapped to the mouse (GRCm38_68) and human (GRCh38) genome reference sequences. Mapping to the mouse genome produced average depth of 27.68-fold (Figure 1). Mapping to the human genome produced an average depth of ∼60 fold in the range chrX:31,081,740-33,507,715 strongly suggesting that this region defines the locus included in the original YAC clone used to make the hDMD transgene and confirming that the region was duplicated based on the coverage depth compared to the mouse genome.

**Figure 1:**
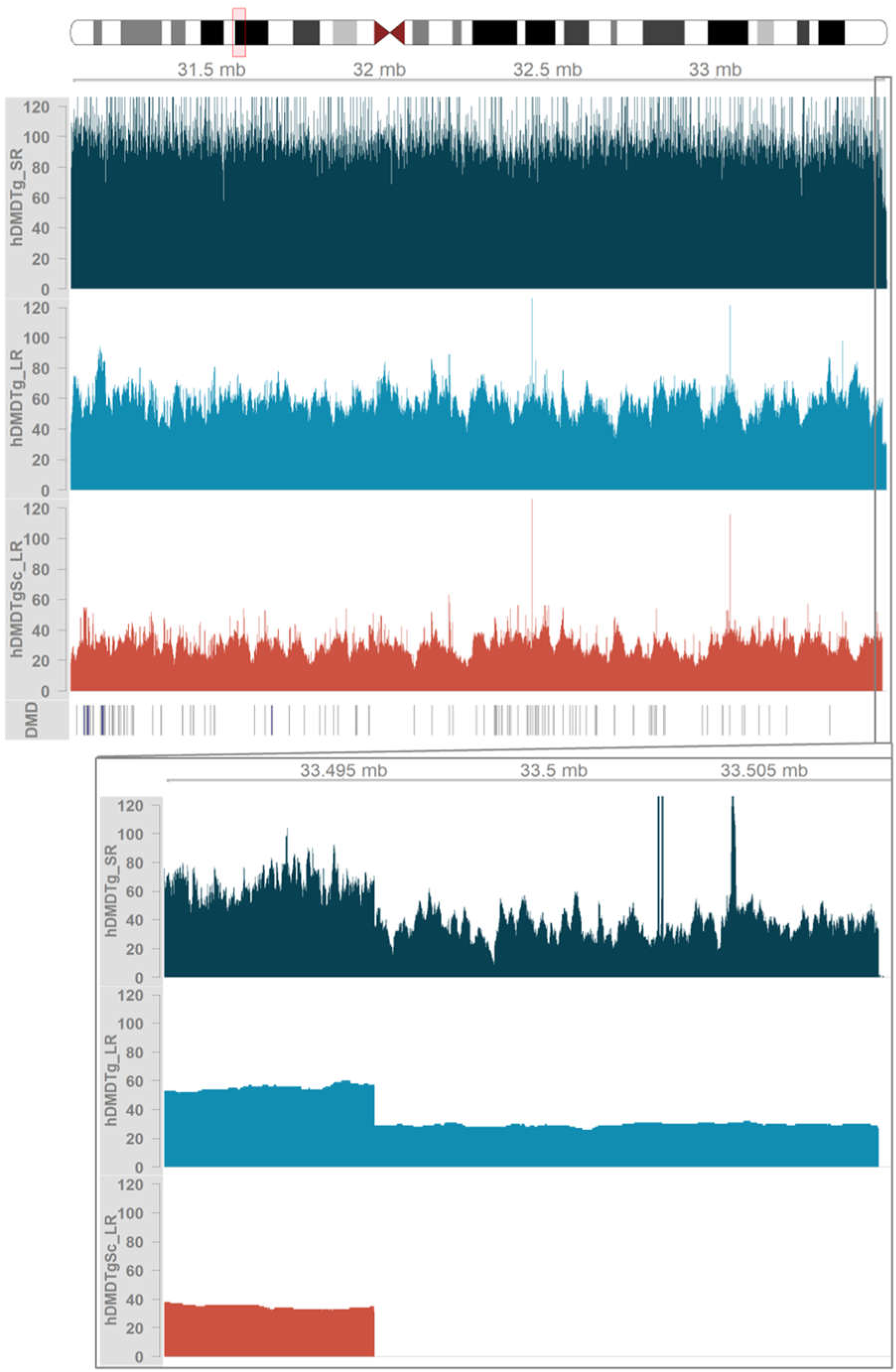
Coverage of the human *DMD* locus in the duplicated (hDMDTg) and single copy (hDMDTgSc) transgenes. Mapping depths at the *DMD* locus for short-read and long-read genomes mapped to the human genome. The red box on the chromosome X ideogram at the top shows the approximate location of the coverage shown. Graphs show the coverage of mapped reads for the short-read (hDMDTg-SR) and long-read (hDMDTg-LR, hDMDTgSc-LR) genomes at this location. The *DMD* exon model is shown below the coverage graphs. At the base of the image a zoomed in section chrX: 33490727-33509715, shows the drop in coverage from position chrX:33,495,727 onwards in the hDMDTg data and no coverage from this position in the hDMDTgSc mice.

To identify the transgene integration site on mouse chromosome 5, we generated a custom mouse reference genome incorporating the *DMD* locus identified from mapping to GRCh38 and the sequence of pYAC4 the transgene vector. Next, we mapped short-read Illumina genome sequencing (GS) data and Nanopore long-read sequence data from the hDMDTg mouse to this custom reference. For short-read GS, 36,781 break ends corresponding to sites of structural variation (deletions, duplications, translocations, or inversions) on chr5 were called with GRIDSS2. In the case of the long read GS, 1769 structural variants (SV) were called on chr5. However, without a non-transgenic littermate control genome, the exact insertion site could not be determined from these investigations alone. There were no translocations between mouse chr5, the human *DMD* locus or the pYAC4 identified from any of these analyses.

De novo assembly of hDMDTg long-read data revealed a collection of related contigs in the hDMDTg assembly graph that corresponded to the transgene (Supplementary Figure 1A). Approximately 10.8 Mb of sequence in the assembly graph which included the *DMD* locus corresponded to yeast chromosomal sequences (Supplementary Figure 1A). Mapping these transgene associated contigs back to the custom GRCm38+*DMD*+pYAC4 reference revealed a breakpoint on chr5: 5:133,626,547 that linked to these inserted transgenic sequences. Re-examination of structural variant calls from both long- and short-read mapped data confirmed the transgene insertion site on mouse chr5, which included a 174 kb duplication of mouse chr 5 (5:g.133,626,547_133,800,724dup) at this location (Figure 2).

**Figure 2:**
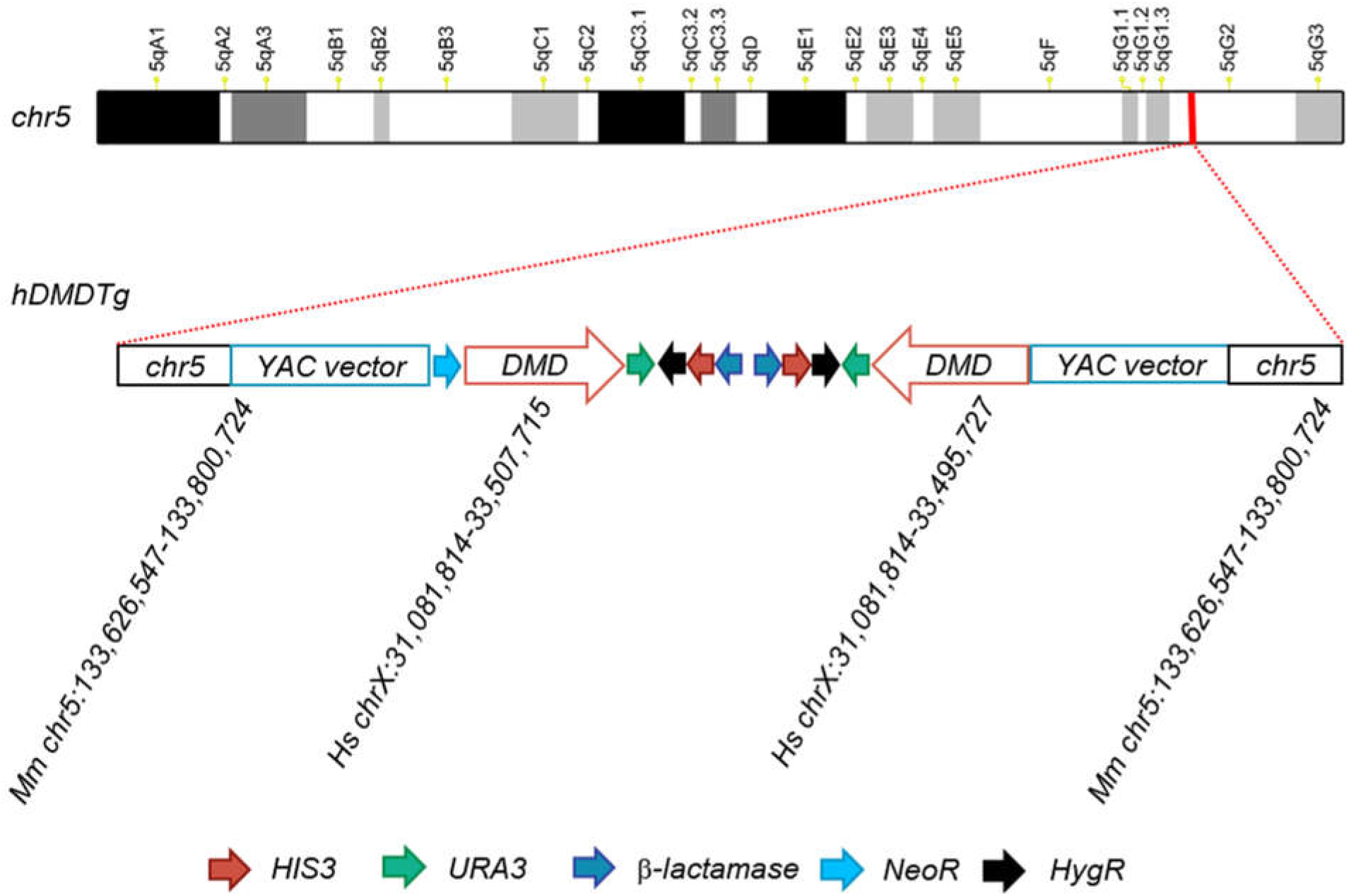
The structure of the duplicated hDMD transgene. Simplified representation of the hDMD duplicated transgene insertion site on mouse chromosome 5 showing the two copies of *DMD* interspersed with YAC vector sequences and the mouse chr 5 duplication.

Comparison of the mapped hDMDTg sequences revealed that the *DMD* loci within the hDMDTg transgene were not exact duplications. One copy ranged from chrX:31,081,740-33,507,715, and the other chrX:31,081,740-33,495,727 (Figure 2). The contig containing *DMD* from the de novo assembly also captured the tail-to-tail orientation of the transgenes consisting of inverted repeats of *URA3, HygR*, *HIS3*, and *β-lactamase* genes (Figure 2). Based on the assembly the NeoR antibiotic resistance cassette is suggested to only occur once at the 5’ end of one of the *hDMD* transgenes (Figure 2).

### Generation of the single copy hDMDTgSc mouse model

To faithfully recapitulate the single-copy nature of the *DMD* gene, we aimed to generate a single-copy hDMD transgenic mouse model (hDMDTgSc) by microinjecting hDMDTg^Tg/0^ embryos with SpCas9 mRNA and gRNAs targeting the NeoR and HygR cassettes on the YAC sequences. This was designed to induce the megabase-scale intervening deletion of only a single transgene. Potential founders were screened by PCR and by identified by amplification of the deletion interval between the NeoR and HygR gRNA target sites, as well as the absence of NeoR amplification. One male founder was identified to carry the desired deletion (Figure 3A).

**Figure 3.**
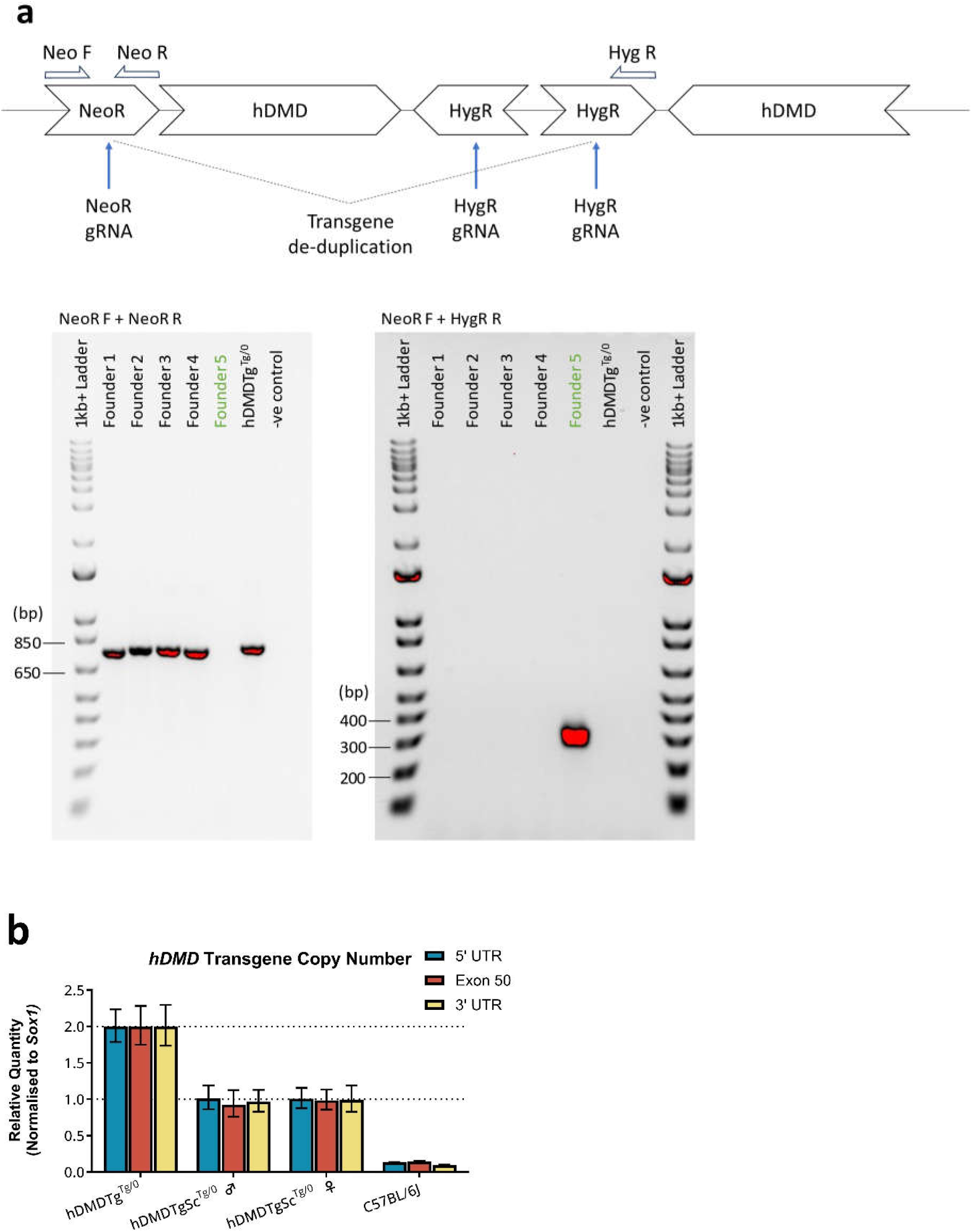
: Genotyping hDMDTgSc mice. (A) Detection of the single transgene deletion in G0 founder mice by PCR. The intervening deletion between the Neomycin resistance (NeoR) and Hygromycin resistance (HygR) cassettes was identified in Founder number 5 from the absence of PCR amplification using primers flanking the NeoR cut site and the presence of amplicon using a forward primer in NeoR and a reverse primer in HygR. The green text indicates the founder with the banding expected from successful transgene deletion. **(B)** Real-time qPCR copy-number determination of the human *DMD* transgene. The assay was performed on genomic DNA extracted from ear notch biopsies of hDMDTg^Tg/0^ mDmd^+/Y^, hDMDTgSc^Tg/0^ mDmd^+/Y^ (male), hDMDTgSc^Tg/0^ mDmd^+/+^ (female) and wildtype C57BL/6J (mDmd^+/Y^) mice. Measurement of three targets, 5’ untranslated region (UTR), exon 50 and 3’ UTR, were normalised against the endogenous mouse *Sox1* gene. Transgene copy number was successfully reduced from two to one in heterozygous hDMDTgSc mice. Error bars indicate the 95% confidence interval of technical replicates.

Following transmission of the de-duplicated transgene to G1 mice, we quantified the hDMD transgene copy number in hDMDTgSc^Tg/0^ mice by qPCR using three sets of primers targeting the 5’ untranslated region (UTR), Exon 50 and 3’ UTR of the *hDMD* transgene. Both G1 male and female hDMDTgSc^Tg/0^ mice were confirmed to carry a single copy of the autosomal *hDMD* transgene, compared to two copies in the hDMDTg^Tg/0^ control (Figure 3B).

### Sequence analysis of hDMDTgSc mice

Mapping Nanopore long-read sequence data from hDMDTgSc mice to our custom reference mouse genome with the human *DMD* locus and pYAC4 sequence, we identified 1462 SVs on chr5. Of all SV called between the hDMDTg and hDMDTgSc data, 196 were concordant, one of which was the 174kb duplication on chr5 containing the transgene insertions site. De novo assembly of hDMDTgSc long-read data also captured the transgene insertion site on mouse chr5; however, a contig corresponding to *DMD* was not assembled from these data by Shasta, and thus, this graph contained only mouse and yeast chromosomal sequences (Supplementary Figure 1B). From comparison to mapped hDMDTg sequences, only the shorter copy of the *DMD* locus (chrX:31,081,740-33,495,727) was retained in the single-copy hDMDTgSc transgene (Figure 1).

### Characterisation of the hDMDTgSc mDmdKO mouse model

To assess if the single copy hDMD transgene in hDMDTgSc mice retains functionality, we characterised male hDMDTg^Tg/0^ mDmd^KO/Y^ and hDMDTgSc^Tg/0^ mDmd^KO/Y^ mice, both strains lacking endogenous mouse dystrophin expression.

Skeletal and heart muscle tissues were collected from p60 mice for mRNA, protein, and histology analysis. We performed quantitative RT-PCR analysis of the dystrophin transcript in quadriceps and heart muscles using a set of primers that detects both the human and mouse transcript by targeting shared sequences. In quadriceps muscle, the expression level of *hDMD* in hDMDTg^Tg/0^ mDmd^KO/Y^ mice is equivalent to that of the *mDmd* in WT C57BL/6J mice, whereas *hDMD* expression in hDMDTgSc^Tg/0^ mDmd^KO/Y^ quadriceps is approximately halved (Figure 4A). In heart muscles, *hDMD* transcript in hDMDTg^Tg/0^ mDmd^KO/Y^ mice is slightly higher than *mDmd* in WT C57BL/6J mice, while *hDMD* expression in hDMDTgSc^Tg/0^ mDmd^KO/Y^ heart is, although somewhat variable, more similar to that of *mDmd* (Figure 4B).

**Figure 4:**
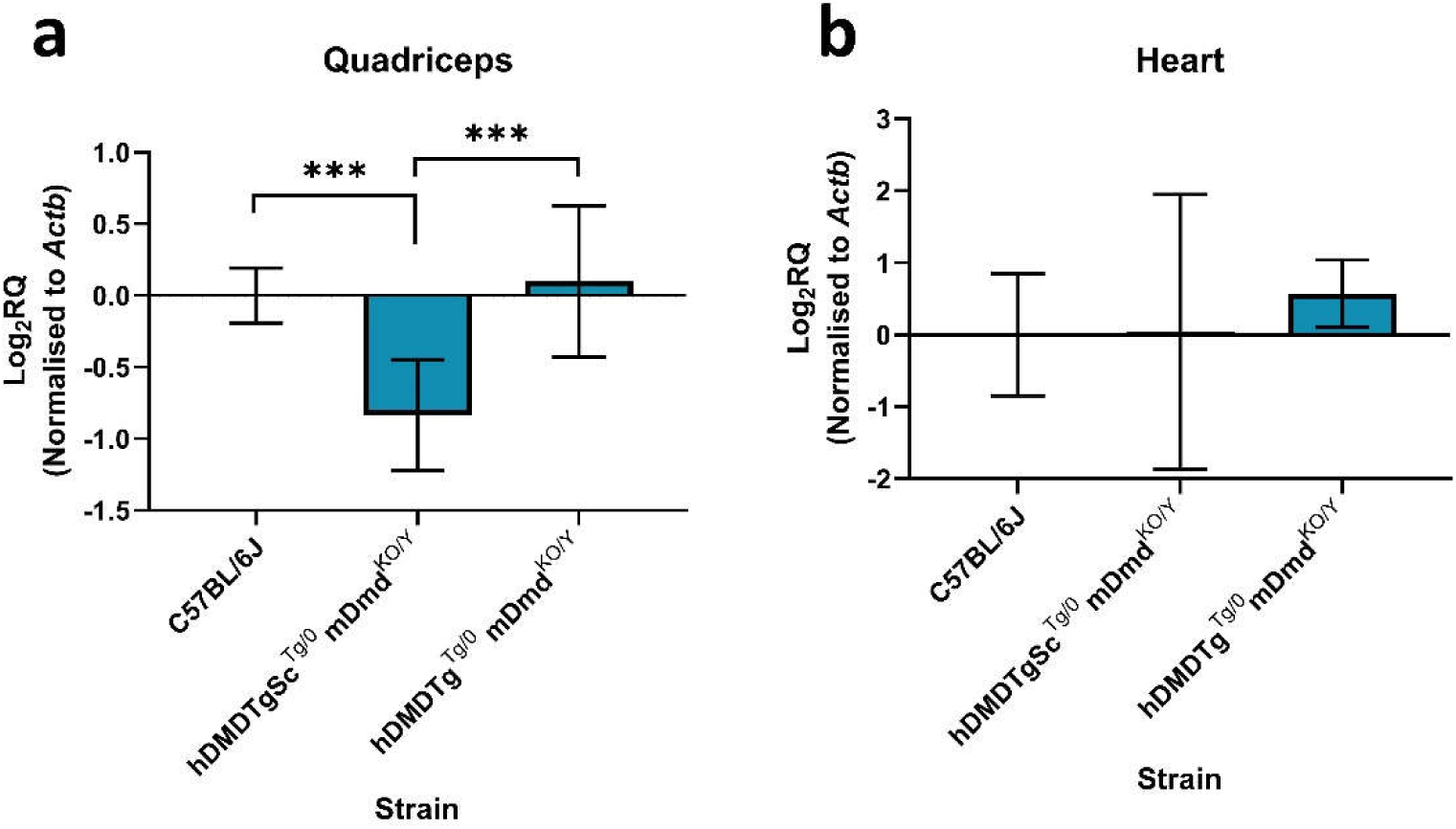
Dystrophin transcript measurement by qRT-PCR. Quantification of (A) skeletal and (B) heart tissues from p60 mice was performed using a set of primers that target both human *DMD* and mouse *Dmd* transcript sequences, normalised against beta-actin (*Actb*). Data presented as log2 relative quantity (RQ) ± 95%CI, one-way ANOVA with Tukey’s multiple comparisons test performed on ddCt values, *** P < 0.001. n=3.

Western blots were performed with the human dystrophin-specific MANDYS106 antibody. Full-length dystrophin protein was detected in tibialis anterior (TA), quadriceps and heart muscles of hDMDTgSc^Tg/0^ mDmd^KO/Y^ mice at a lower level compared to hDMDTg^Tg/0^ mDmd^KO/Y^ mice (Figure 5). The quadriceps western blot was re-probed with MANDYS8, which is cross-reactive with both human and mouse dystrophin, confirming that MANDYS106 only detects human dystrophin.

**Figure 5:**
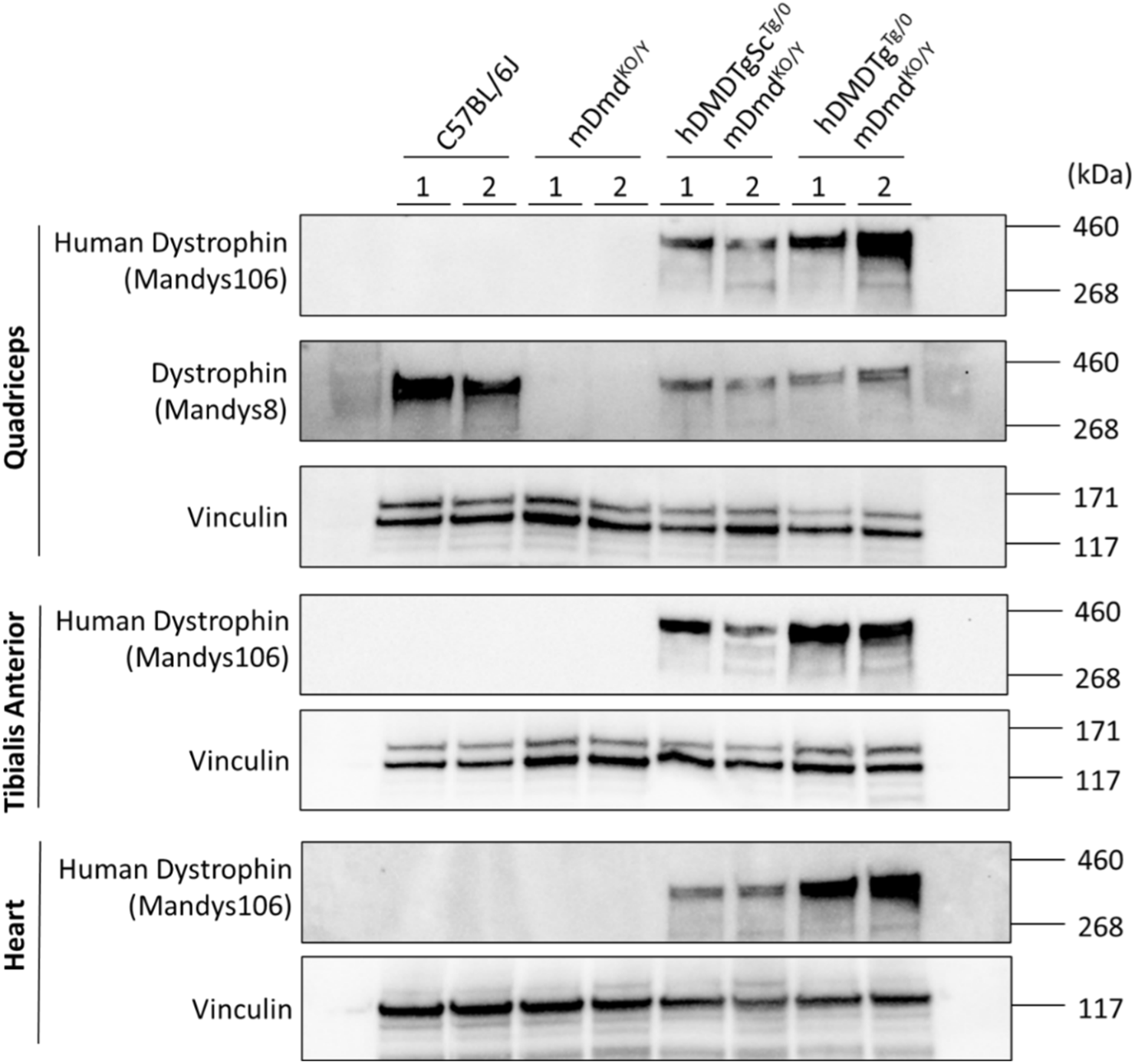
Detection of human dystrophin protein in muscle tissue by western blot. Analysis was performed on skeletal hindlimb muscles, tibialis anterior and quadriceps, as well as heart muscle protein lysates from p60 mice using the human-specific antibody against dystrophin MANDYS106. The tibialis anterior blot was also re-probed with the cross-reactive antibody MANDYS8 to demonstrate the species-specificity of MANDYS106. Vinculin was probed as the loading control. Human dystrophin protein expression in hDMDTgSc^Tg/0^ mDmd^KO/Y^ mice is reduced in all three tissues compared to hDMDTg^Tg/0^ mDmd^KO/Y^ mice.

Immunofluorescence staining of dystrophin with MANDYS106 in TA, quadriceps and heart muscles showed the expected sarcolemmal localisation of the human dystrophin protein in hDMDTgSc^Tg/0^ mDmd^KO/Y^ muscle, as for hDMDTg^Tg/0^ mDmd^KO/Y^ muscle tissue (Figure 6, Supplementary Figure 2).

**Figure 6:**
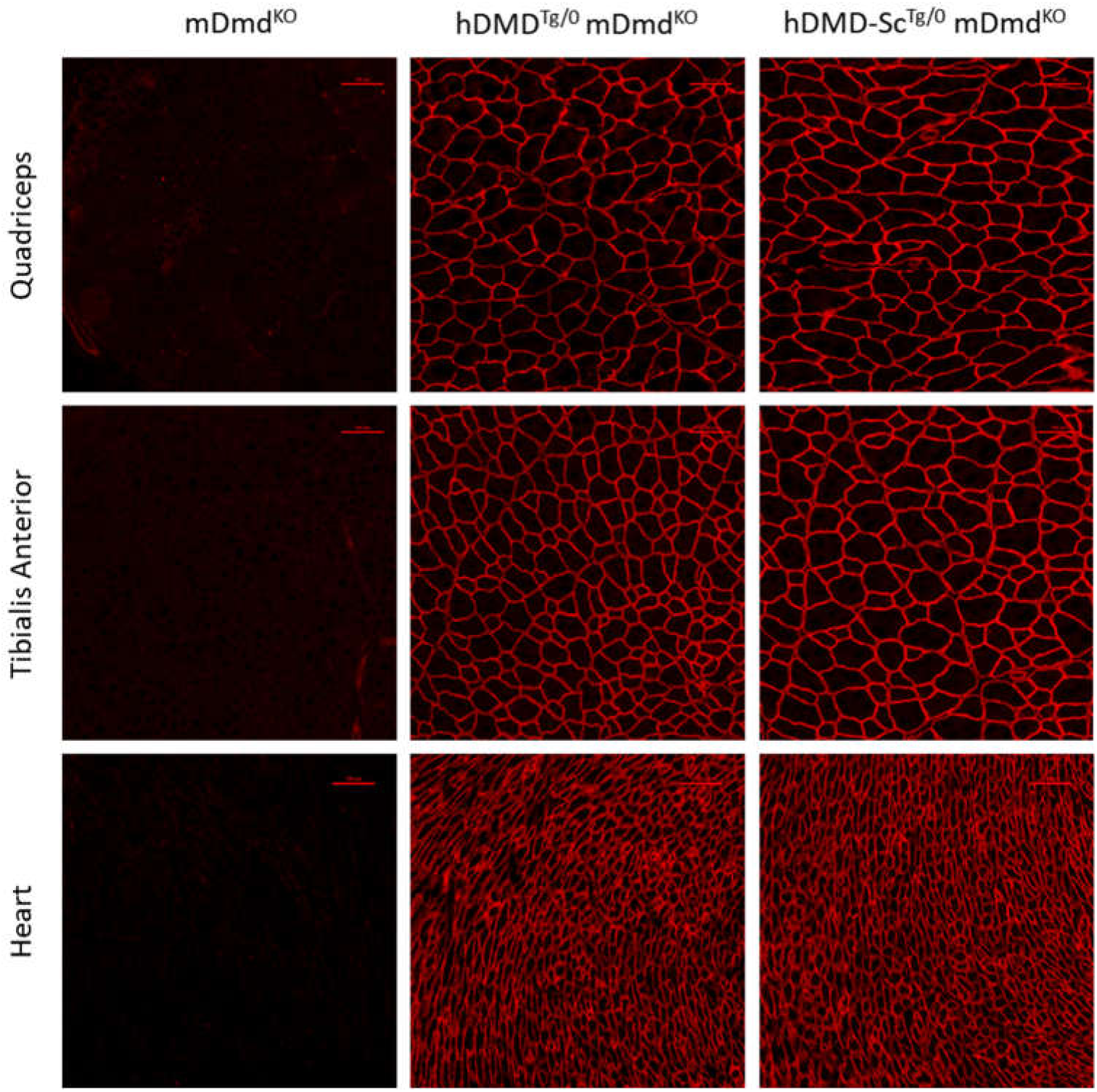
Immunofluorescence localisation of the human dystrophin protein. Immunostaining was performed on tibialis anterior, quadriceps and heart muscle tissue sections from p60 mice using the MANDYS106 primary antibody. Scale bar 100 µm.

Histological analysis of hDMDTgSc^Tg/0^ mDmd^KO/Y^ muscle sections stained with hematoxylin and eosin (H&E) revealed normal muscle histology with nuclei localised at the sarcolemmal region, consistent muscle fibre size, and no evidence of damage or infiltrating cells (Figure 7). Conversely, mDmd^KO/Y^ muscle was dystrophic, with many centralised nuclei indicating muscle regeneration as well as the presence of cellular infiltrates.

**Figure 7:**
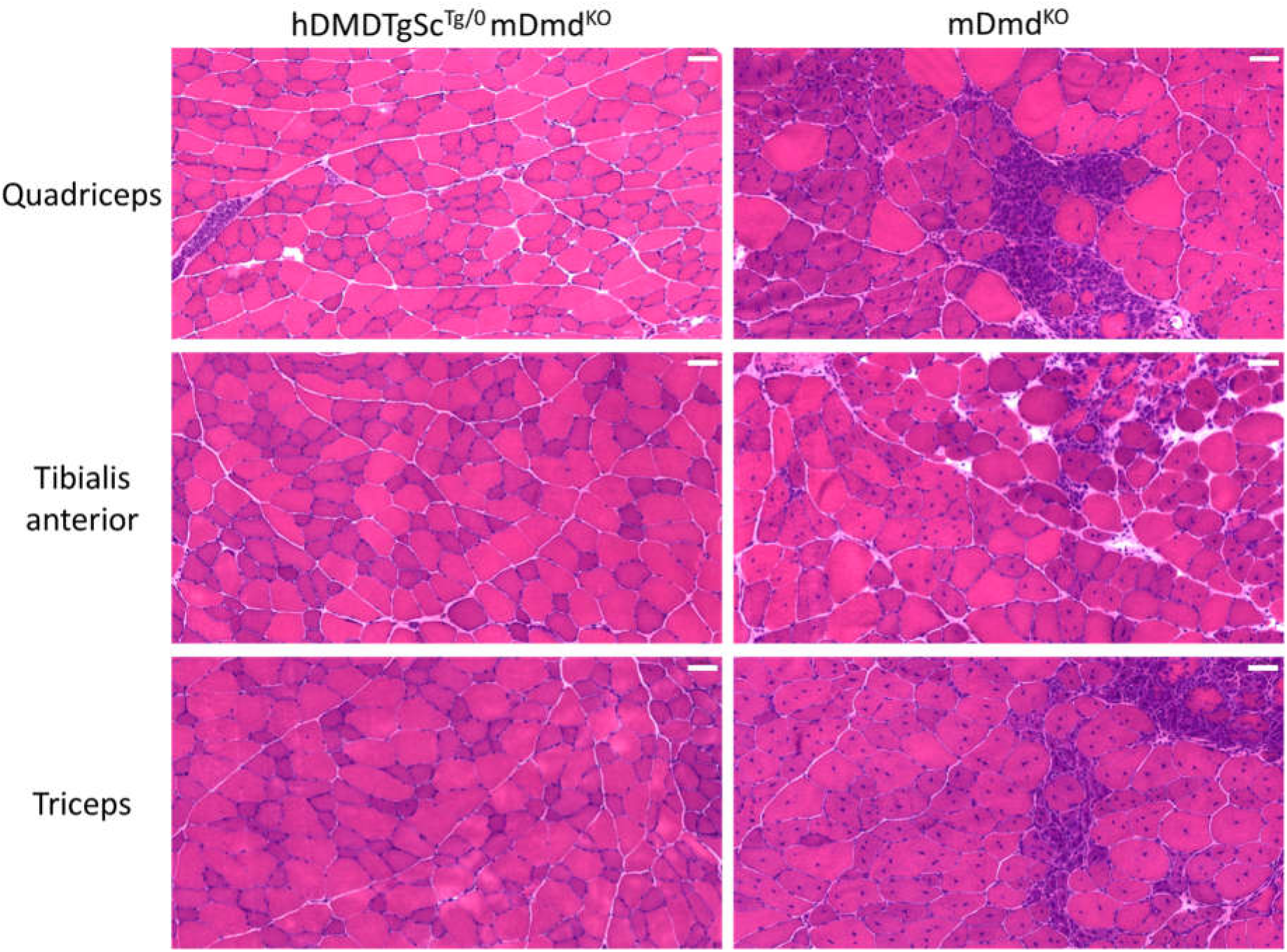
Histological analysis of hDMDTg^Tg/0^ mDmd^KO/Y^ and mDmd^KO/Y^ mice. Hematoxylin and eosin staining was performed on quadriceps, tibialis anterior and triceps muscle tissue sections from p60 mice. Scale bar 50 µm.

We performed forelimb grip strength testing followed by serum collection for creatine kinase analysis on mice at postnatal day 32 (p32) and day 60 (p60). Forelimb grip strength testing showed that hDMDTgSc^Tg/0^ mDmd^KO/Y^ mice have similar maximum and mean grip strength to hDMDTg^Tg/0^ mDmd^KO/Y^ mice at p32 and that both these strains have no significant difference in strength compared to wildtype C57BL/6J control mice, demonstrating functional rescue of the reduced grip strength caused by the endogenous *mDmd* knock-out (Figure 8A,B). hDMDTgSc^Tg/0^ mDmd^KO/Y^ and hDMDTg^Tg/0^ mDmd^KO/Y^ also do not experience the same onset of fatigue caused by repeated measurements as observed in mDmd^KO/Y^ mice (Figure 8C,D).

**Figure 8:**
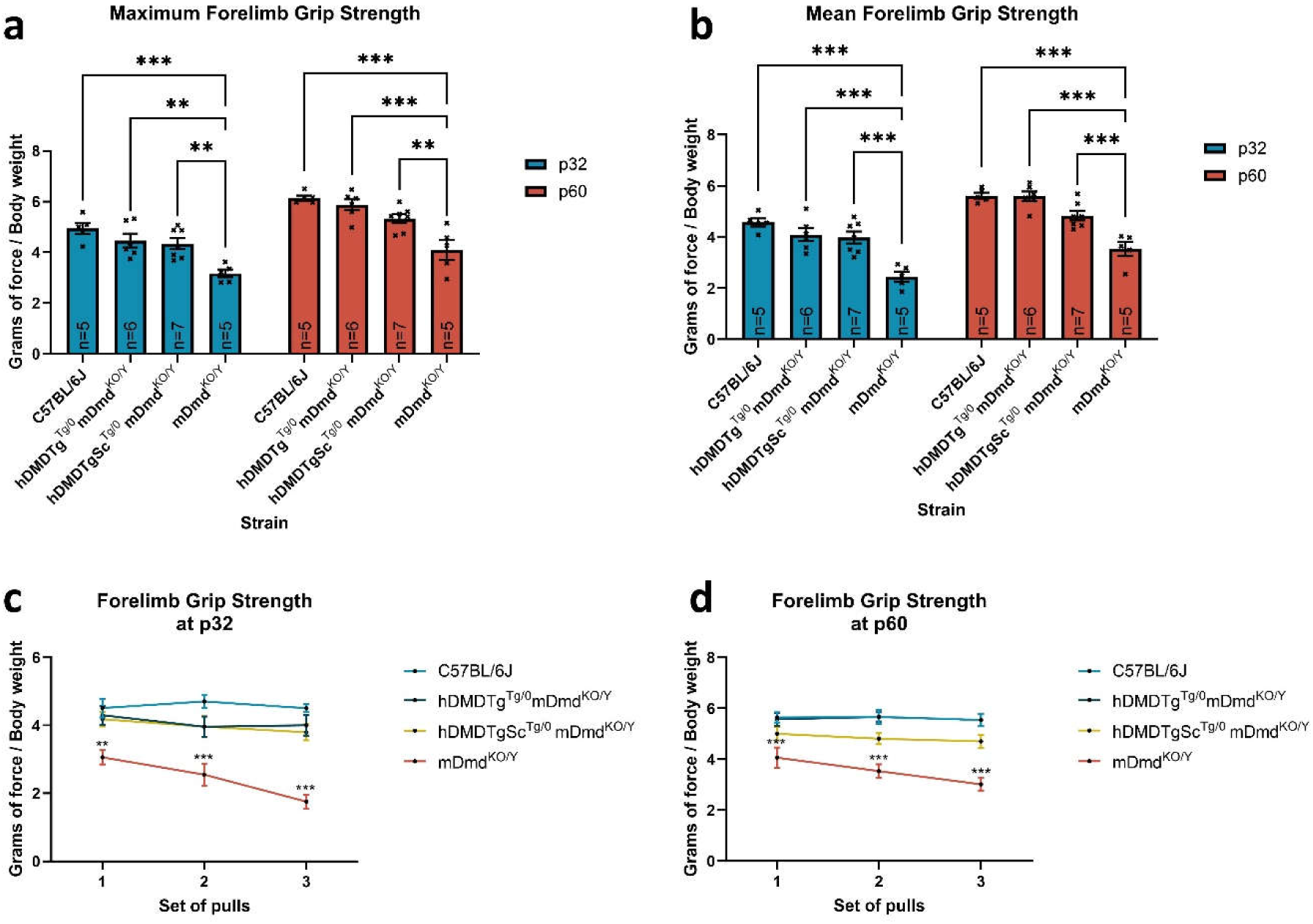
Forelimb grip strength testing to measure muscle function and performance of mice. Strength (grams of force) of hDMDTg^Tg/0^ mDmd^KO^ and hDMDTgSc^Tg/0^ mDmd^KO^ mice were compared against WT C57BL/6J and mDmdΔEx51 (mDmd^KO/Y^) controls, normalised to body weight at p32 and p60. (A) Maximum force is the highest recorded measure among three sets of five pulls, and (B) mean force is the average of the highest recorded measures of each of the three sets. Data presented as mean ± SEM, two-way repeat measures ANOVA with Tukey’s multiple comparisons test ** P < 0.01, *** P < 0.001. Samples sizes (n) indicated in within bars in graph. (C,D) Averages of the highest recorded measures for each set of pulls, presented as mean ± SEM, two-way repeat measures ANOVA, Dunnett’s multiple comparisons test with C57BL/6J as control group, ** P < 0.01, *** P < 0.001.

The serum samples collected after the forelimb grip strength testing were used to assess serum creatine kinase (CK) levels, an indirect measure of muscle damage. Both hDMDTg^Tg/0^ mDmd^KO/Y^ and hDMDTgSc^Tg/0^ mDmd^KO/Y^ mice have a similar reduction of serum CK levels to WT C57BL/6J controls at p32 and p60 (Figure 9).

**Figure 9:**
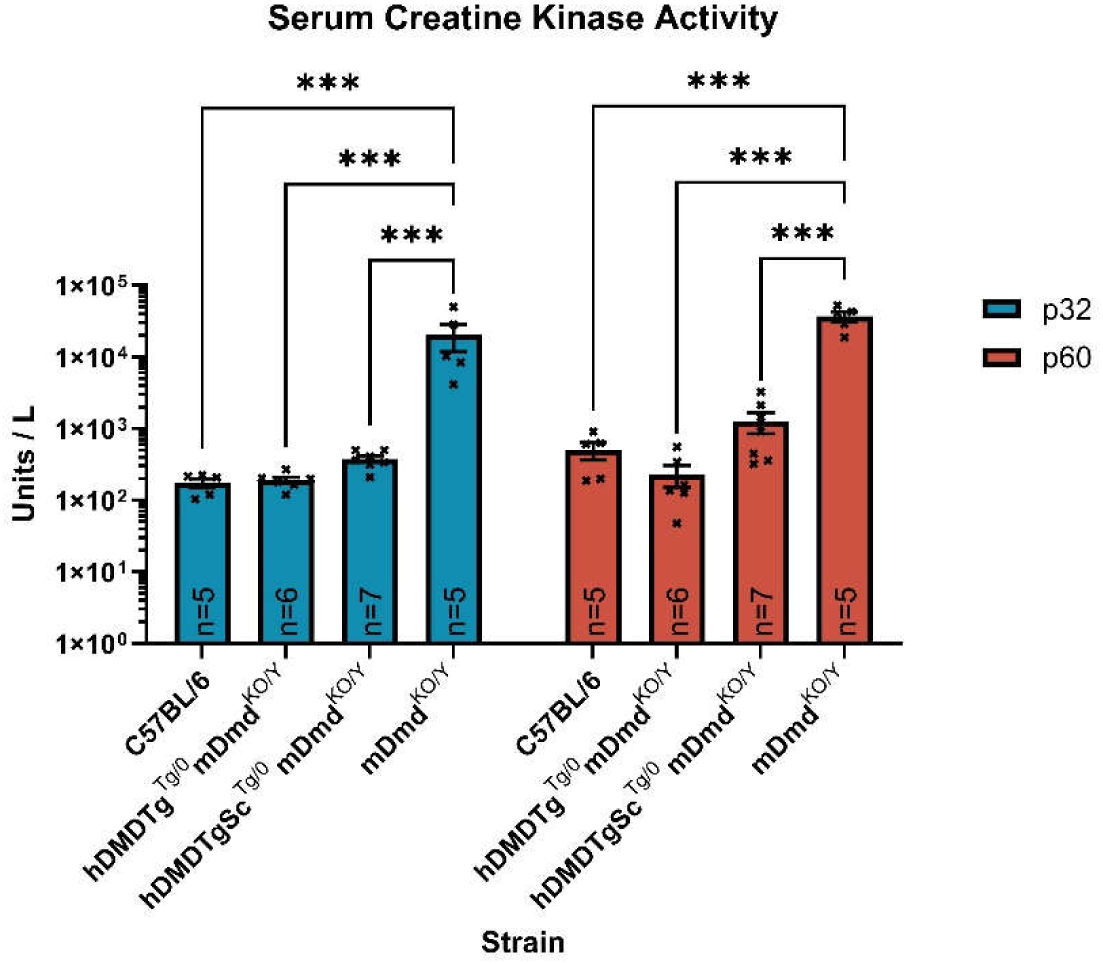
Serum creatine kinase (CK) biomarker level measurement of mice. Serum CK was assayed as an indirect measure of muscle damage and sarcolemma leakage at p32 and p60. The single-copy transgene in hDMDTgSc^Tg/0^ mDmd^KO/Y^ mice rescues the elevated serum CK caused by endogenous *mDmd* knock-out to levels that are not significantly different than either WT C57BL/6J or hDMDTg^Tg/0^ mDmd^KO/Y^ mice. Data presented as mean ± SEM, two-way repeated measures ANOVA with Tukey’s multiple comparisons test *** P < 0.001. Samples sizes (n) indicated in first graph.

## Discussion

Validated humanised animal models are important tools for pre-clinical evaluation of genetic therapies as they permit in vivo testing of sequence-specific reagents that can be directly translated into patients. Generation of a hDMDTg mouse model containing the entire *hDMD* locus randomly integrated into mouse chromosome 5 was a significant advance, particularly given the size of the transgene (2.4 Mb). However, it was subsequently shown that two copies of the transgene had inserted, which does not accurately reflect the copy number of the *DMD* gene [14]. Furthermore, derivatives of the hDMDTg line such as the del52 exon deletion disease model are more difficult to engineer and validate. While these double-copy humanised models can be used for the testing of ASOs [15], they are not ideal for evaluating strategies that are reliant on DNA double-stranded breaks such as CRISPR-Cas9 gene editing [16]. This is because large intervening deletions between the target sites of the two transgene copies can complicate the analysis of DNA repair outcomes.

Here, we describe the generation and characterisation of the hDMDTgSc mouse model carrying a single functional copy of the 2.4 Mb *hDMD* transgene. Large megabase-scale DNA modifications have been previously demonstrated to be achievable through CRISPR microinjection[17–20]. We performed a detailed genetic characterisation of the original duplicated hDMDTg transgene structure by analysis of Illumina short-read and Nanopore long-read sequencing data to determine unique targetable regions. Using gRNAs that flank one copy of the transgene, we readily achieved transgene de-duplication of the hDMDTg mouse model by CRISPR zygote microinjection and demonstrated stable germline transmission.

We also successfully resolved the exact YAC integration site in chromosome 5 and can confirm that no genes were disrupted at the integration site. Long-read sequencing helped us confirm that a single copy of the hDMD transgene remained intact in hDMDTgSc mice following the de-duplication process, and its reduced copy number was also reflected in the reduced mapping depths from sequencing.

There are some limitations to the genetic analyses we performed via next-generation sequencing. Using only nanopore long-read sequences, we were not able to unambiguously assemble the full-length hDMDTgSc and hDMDTg transgenes despite achieving a total assembly length similar to the length of the mm10 genome reference and a relatively high contig N50 (Supp Table 1). The exact assembly and ordering of the yeast chromosomal sequences that flank the DMD loci in the duplicated and single-copy transgenes remains uncertain. Additional long-read sequencing with high-quality PacBio circular consensus sequences would likely be sufficient to create a near-continuous assembly of the transgenic mouse genome as has been done for the human T2T reference sequence [21].

From phenotypic, biochemical, molecular, and histological characterisation of hemizygous hDMDTgSc mice lacking endogenous mouse dystrophin, we found that the single hDMD transgene is sufficient to rescue the dystrophic phenotype observed in mDmd knock-out mice at a similar level to the hDMDTg mDmdKO strain. This is despite the reduced transgene transcription in skeletal muscle driven by the human promoter sequence, which is likely not optimally regulated by the mouse transcription machinery [22]. Additionally, there appears to be a slight reduction of hDMDTgSc mDmdKO mouse strength compared to hDMDTg mDmdKO mice at p60 that might reflect a high but incomplete level of functional rescue-more studies are needed to determine if it is due to transgene expression not being as high as in hDMDTg mDmdKO mice or if it could be attributed to experimental variation.

Our hDMDTgSc mouse model has considerable potential for the pre-clinical testing of sequence-specific ASOs or CRISPR therapeutics as, in principle, any causative DMD mutation can be engineered into the single-copy 2.4 Mb *hDMD* transgene, such as models lacking *DMD* exons 45 or 52 like those previously generated from the double transgenic mice. This enables ready generation of humanised DMD disease mouse models, which has an advantage over the more labour-intensive and inefficient partial sequence humanisation methods where exon knock-ins must be performed for each separate exonic target [23]. Crossing these human transgene knock-out mice with mice lacking endogenous dystrophin expression would result in a disease model suitable for the testing of CRISPR therapy strategies, and could also be inter-crossed with a mouse strain lacking both endogenous dystrophin and utrophin expression to simulate an even more severe dystrophic phenotype [24].

## Materials and Methods

### Animal studies

All procedures and experiments using mice were approved by the South Australian Health and Medical Research Institute (SAHMRI) Animal Ethics Committee and conducted per the Australian code for the care and use of animals for scientific purposes.

Mice were housed in 13.5-hour light/10.5-hour dark cycles and were fed the Teklad Global 2918 rodent diet *Ad Libitum*. Except where indicated, male mice were used in characterisation experiments. Tg(DMD)72Thoen/J (hDMDTg) were obtained from JAX (stock #018900) and maintained in-house [8]. mDmdΔEx51 (mDmd^KO^) and hDMDTgSc were generated as described in the sections below and maintained in-house. We maintain hDMDTgSc mice as a tick-over colony of homozygotes. C57Bl/6J mice are maintained in house (JAX stock #000664).

### Mouse generation

To create the intervening deletion of a single copy of the *hDMD* transgene, we designed SpCas9 gRNAs targeting the Neomycin and Hygromycin antibiotic resistance cassettes in the integrated YAC sequence. Complementary gRNA oligos with overhangs facilitating golden gate assembly were phospho-annealed and inserted into pSpCas9(BB)-2A-Puro (PX459) V2.0 [25]. pSpCas9(BB)-2A-Puro (PX459) V2.0 was a gift from Feng Zhang (Addgene plasmid # 62988 ; http://n2t.net/addgene:62988 ; RRID:Addgene_62988). Following plasmid validation by *BbsI* diagnostic digest and Sanger sequencing (Australian Genome Research Facility), gRNAs were amplified using NEB Phusion in HF buffer with primers containing a 5’ T7 promoter sequence. PCR amplicons purified using the QIAquick PCR Purification Kit (Qiagen) were used to generate gRNA using the HiScribe T7 Quick High Yield RNA Synthesis Kit (NEB) and purified using the RNeasy Mini Kit (Qiagen).

hDMDTg^Tg/Tg^ mouse sperm was used for *in-vitro* fertilisation of C57BL/6J oocytes, and 2PN embryos were injected with a microinjection mix containing a final concentration of 100 ng/μL SpCas9 mRNA and 50 ng/μL each of NeoR and HygR gRNA. This was followed by oviduct transfer into pseudo-pregnant females. Intervening transgene deletion was detected in founders by PCR amplification using the forward NeoR and reverse HygR primers.

mDmd^KO^ mice were generated by deletion of *Dmd* Exon 51 via CRISPR microinjection of C57BL/6J embryos (manuscript submitted).

### DNA PCR and Sanger sequencing

Genomic DNA (gDNA) was isolated from mouse ear punch biopsies using the PCR Template Preparation kit (Roche). Targets was PCR amplified using Taq Polymerase (Roche) and FailSafe™ Buffer J (Epicentre) and resolved on a 1% agarose TBE gel.

To prepare samples for sequencing, PCR reactions were cleaned up using the QIAquick PCR Purification kit (Qiagen). Samples with sequencing primers added were sent to the Australian Genome Research Facility (AGRF) for the purified DNA Sanger sequencing service.

### Short-read genome sequencing

Library prep and sequencing was performed as a service by the Australian Genome Research Facility (AGRF). Genomic DNA extracted from hDMDTg mice liver were sheared and PCR-free sequencing libraries prepared. From these libraries, 150 base pair, paired-end reads were generated on a NovaSeq 6000. Image analysis was performed in real time by the NovaSeq Control Software (NCS) v1.7.5 and Real Time Analysis (RTA) v3.4.4. Then the Illumina bcl2fastq v2.20.0.422 pipeline was used to generate the final fastq files.

#### Mapping

All mapping of short read data was carried out using BWA-MEM v 0.7.17 [26] setting option -K, the number of input bases, to 100,000,000 per batch and all other options as default.

A custom reference genome was prepared which combined the GRCm38.p3 mouse genome (GenBank: GCA_000001635.5), a segment of human chromosome X (NCBI RefSeq: NC000023.11; chrX:31,081,740-33,507,715) and pYAC4 (GenBank: U01086.1). The chrX segment was defined by identifying the extents of reads accurately mapped to the region flanking *DMD* on chrX in the hs38DH human genome (created using the run-gen-ref script from bwakit) [27]. The pYAC4 vector sequence was used as a proxy for the pRANT-11 and pLGTel-1 vectors that were originally used for cloning of the hDMD YAC transgene and for which no sequence data are available [28].

Alignments were sorted by genome coordinate positions using samtools sort v 1.17 and duplicate reads were marked using sambamba markdup v 0.8.2 [29].

#### Variant calling

Structural variants were called using GRIDSS v 2.12.0 with default settings [30].

### Oxford nanopore genome sequencing

High molecular weight genomic DNA was extracted from homozygous hDMDTg and hDMDTgSc mice liver using the Monarch HMW DNA Extraction Kit for Tissue (NEB). Oxford Nanopore Technologies Promethion genome sequencing was performed as a service by Novogene, (Novogene HK). Sequence libraries were prepared using the DNA ligation sequencing protocol with the SQK-LSK-110 kit.

Sequence data were generated using a Promethion 9.4.1 flow cell. Bases were called with bonito v0.6.2 (Oxford Nanopore Technologies) with the dna_r9.4.1_e8_sup@v3.3 super accurate base calling model.

#### Mapping

Oxford Nanopore long-read data were mapped to the custom GRCm38+*DMD*+pYAC4 genome described above and also the human genome reference GRCh38 using minimap2 v2.17-r941 with default parameters for nanopore sequencing.

#### Variant calling

Structural variants were called from mapped long read genome sequencing with Sniffles2 using default settings [31].

#### Assembly

Nanopore sequence reads were assembled de novo using Shasta v 0.11.1 [32] using the Nanopore-May2022 model adjusting the minimum length of input reads to 5 kb. To identify contigs associated with the hDMD transgene sequence, the Assembly.fasta outputs were mapped to the custom GRCm38+*DMD*+pYAC4 genome described above with minimap2 and assembled into super contigs using RagTag v 2.1.0 [33]. The contigs from the Assembly.fasta files were also converted into a BLAST v 2.14.1+ library that was queried with 500 bp segments from the 5’ and 3’limits of the *DMD* locus in the hDMDTgSc transgene. Assembly graphs were viewed using Bandage v 0.8.1.

### Transgene copy number determination

qPCRs were performed to quantify the *hDMD* transgene copy number and were done in technical triplicate. Each reaction well was set up using SYBR™ Green PCR Master Mix (Applied Biosystems), 50 nM of each forward and reverse primer and 10 ng gDNA in a 15 μL reaction volume. qPCR data was normalised to endogenous *Sox1.* Reactions were run on the QuantStudio™ 3 Real-Time PCR System (Applied Biosystems) using the Fast Run Mode with the following reaction conditions: 95°C for 20 s, followed by 40 cycles of 95°C for 1 s and 60°C for 20 s, and a dissociation protocol to obtain a melt curve for all samples running from 60°C to 95°C.

### RNA extraction and qRT-PCR

Muscle tissues were bead homogenised in 500 µL TRIzol reagent (Invitrogen) using Lysing Matrix D (MP Biomedicals). 100 µL of chloroform was added and samples were shaken vigorously, incubated at RT for 3 mins then centrifuged at 10 000 g for 18 min at 4°C to separate phases. The aqueous phase was isolated, mixed with an equal volume of ethanol, and spun through a RNeasy column from the RNeasy Mini kit (Qiagen) at 8 000 g for 30 s. The column was washed with 700 µL buffer RW1 and twice with 500 µL buffer RPE. RNA was eluted in 30 µL of nuclease-free water, quantified by nanodrop and stored at -80°C. cDNA was generated from 1 µg RNA using the High-Capacity RNA-to-cDNA kit (Applied Biosystems) as per manufacturer’s protocol.

qRT-PCRs were performed to quantify dystrophin transcript expression in technical and biological triplicate. The primers used were designed to target shared sequences in both *hDMD* and *mDmd* transcript. Each reaction well was set up using SYBR™ Green PCR Master Mix (Applied Biosystems), 50 nM of each forward and reverse primer and 1 µL of cDNA in a 15 μL reaction volume. qPCR data was normalised to beta-actin (*Actb*) expression. Reactions were run on the QuantStudio™ 3 Real-Time PCR System (Applied Biosystems) with the same parameters listed in the transgene copy-number determination section above. Analysis was performed using the Design and Analysis Software 2.7.0. Transcript expression data presented as log2(RQ) to show proportional changes.

### Western blot

Tissues were bead homogenised in disruption buffer (75 mM Tris HCl pH 6.8, 15% SDS, 1x cOmplete Protease Inhibitor Cocktail EDTA (Roche)), using Lysing Matrix D (MP Biomedicals). 1/10 diluted lysates were quantified using the Pierce BCA protein assay (Thermo Scientific). 30 μg of protein lysates were mixed with equal parts 2x Loading buffer (21% w/v glycerol, 0.001% w/v bromophenol blue, 5% β-Mercaptoethanol), boiled at 95°C for 5 mins and spun at 21 300 rcf for 5 mins prior to loading in NuPAGE Tris-Acetate gels (Invitrogen). Gels were run at 150 V for 1 h and transferred onto 0.2 µm PVDF membranes using the High MW setting on the Trans-Blot Turbo (Bio-Rad). Membranes were blocked with 10% skim milk in TBS-T for 2h with gentle agitation at RT. Dystrophin was probed with 1:500 Anti-Dystrophin antibody, clone 2C6 MANDYS106 (Sigma-Aldrich) in 2% skim milk in TBS-T overnight at 4°C. The following day, membranes were washed three times for 5 mins with TBS-T and incubated with 1:5 000 secondary anti-mouse antibody in 2% skim milk in TBS-T for 1h at RT. After another set of TBS-T washes, membranes were developed with SuperSignal West Pico Plus Chemiluminescent Substrate (Thermo Scientific), and chemiluminescent detection was performed on the ChemiDoc MP system (Bio-Rad) using automated exposure settings for intense bands. Membranes were stripped with Restore PLUS Western Blot Stripping Buffer (Thermo Scientific) prior to re-probing. All dystrophin was probed with 1:500 anti-dystrophin antibody MANDYS8. Vinculin was probed with 1:1 000 Anti-vinculin antibody, V9131 (Sigma-Aldrich).

### Immunofluorescence

Fresh muscle was snap-frozen in dry-ice-cooled isopentane and immediately embedded in Tissue-Tek O.C.T. 10 μm thick cryostat-sliced transverse sections were permeabilised with PBS + 0.3% Triton X-100 for 10 minutes, blocked with 10% FCS in PBS + 0.3% Triton X-100 for 1 h at RT, and treated with 1x ReadyProbes™ Mouse-on-Mouse IgG Blocking Solution (Invitrogen) as per manufacturer instructions. Slides were incubated with 1:100 Anti-Dystrophin antibody, clone 2C6 MANDYS106 (Sigma-Aldrich) diluted in blocking solution overnight at 4°C in a humidified chamber. The next day slides were washed with PBS for 3x 5mins and incubated with secondary anti-mouse antibody conjugated with Alexa Fluor® 594 for 1h at RT. Slides were washed with PBS for 3x 5mins and mounted with ProLong™ Gold Antifade Mountant with DAPI (Invitrogen) and coverslip. Images were taken on a Nikon Eclipse Ti2 microscope with a DS-Qi2 camera. The same image processing was applied to images from the same muscle group.

### Histological analysis of muscles

Fresh muscle was mounted on 10% gum tragacanth on a piece of cork, snap frozen in liquid-nitrogen-cooled isopentane and immediately stored at -80°C. For skeletal muscles, 10 μm thick cryostat-sliced transverse sections were stained with Lillie Mayer’s Hematoxylin and Eosin, as described by Wang et al. (2017) with the following modifications to the dehydration steps: 2 dips of 70% ethanol, 4 dips of 95% ethanol, 2x 1 min of 100 ethanol and 2x 1 min of xylene. For heart muscles, 10 μm thick cryostat-sliced transverse sections were stained with Lillie Mayer’s Hematoxylin for 6 mins, rinsed with running tap water for 1min, destained with 0.5% acid alcohol and developed in Scott’s tap water for 1 minute [34]. After rinsing with running water for 2 minutes, slides were stained with eosin for 3 mins and quickly rinsed with running tap water before dehydrating through an increasing concentration of ethanol and xylene as described in the skeletal muscle protocol. Slides were mounted with DPX mountant (Sigma-Aldrich) and coverslip. Images were taken using the 3DHISTECH Pannoramic 250 Flash Slide Scanner at 40x magnification. The same image processing was applied to all images.

### Forelimb grip strength test

Muscle strength measurement was performed by a forelimb grip strength behaviour task [35]. Mice were tested at p32 and p60 using a grid attached to a Centor Easy 10N (Andilog) force gauge. Mice were suspended by the tail and allowed to grasp the grid tightly with both front paws, then pulled along a straight horizontal line away from the force gauge until their grip on the grid was broken.

Measurements were repeated 4 times per set, and 3 sets were performed with a 1-minute rest interval between sets. The highest measured rep was recorded for each set and normalised against body weight. Max force reported is the highest measured rep among all sets, and mean force reported is the average of the highest measured reps for each of all 3 sets.

### Serum creatine kinase measurement

30 mins after grip strength testing, Emla 5% local anaesthetic cream was applied to the mice’s tails, and they were warmed in a Mini Thermacage (datesand) for 10 mins at 36°C to dilate the tail vein. Blood samples were collected in microvettes from a nick to the lateral tail vein. The blood was allowed to clot at room temperature for 30 mins, and the clot was separated from the serum by centrifuging at 2 000 g for 10 mins at 4°C and immediately stored at -80°C. Mouse serum creatine kinase (CK) levels were measured by Gribbles Veterinary Pathology using the ADVIA Creatine Kinase CK_L assay on the ADVIA 1800 Chemistry System.

## Availability of data and materials

The sequencing dataset supporting the conclusions of this article is available in the NCBI Sequence Read Archive, [unique persistent identifier and hyperlink to dataset(s) in http:// format].

## Acknowledgements

Research was performed at SAHMRI on Kaurna Country. We pay respects to the Kaurna people of the Adelaide Plains and to all Aboriginal and Torres Strait Islander people. SAHMRI is committed to embracing knowledge and culture as we continue our working journey to incorporate Aboriginal health research across all our themes and further reconciliation.

This research is supported by the SAHMRI core facilities: Bioresources, Light Microscopy and Laser Capture, Histology and Cryogenics. Generation of mouse strains was carried out with the assistance of the South Australian Genome Editing (SAGE) Facility, the University of Adelaide and the South Australian Health and Medical Research Institute (SAHMRI). SAGE is supported by Phenomics Australia. Phenomics Australia is supported by the Australian Government through the National Collaborative Research Infrastructure Strategy (NCRIS) program. We would also like to thank Melissa White for her comments and suggestions in writing this paper.

## Supplementary materials

**Supplemental Table 1:**
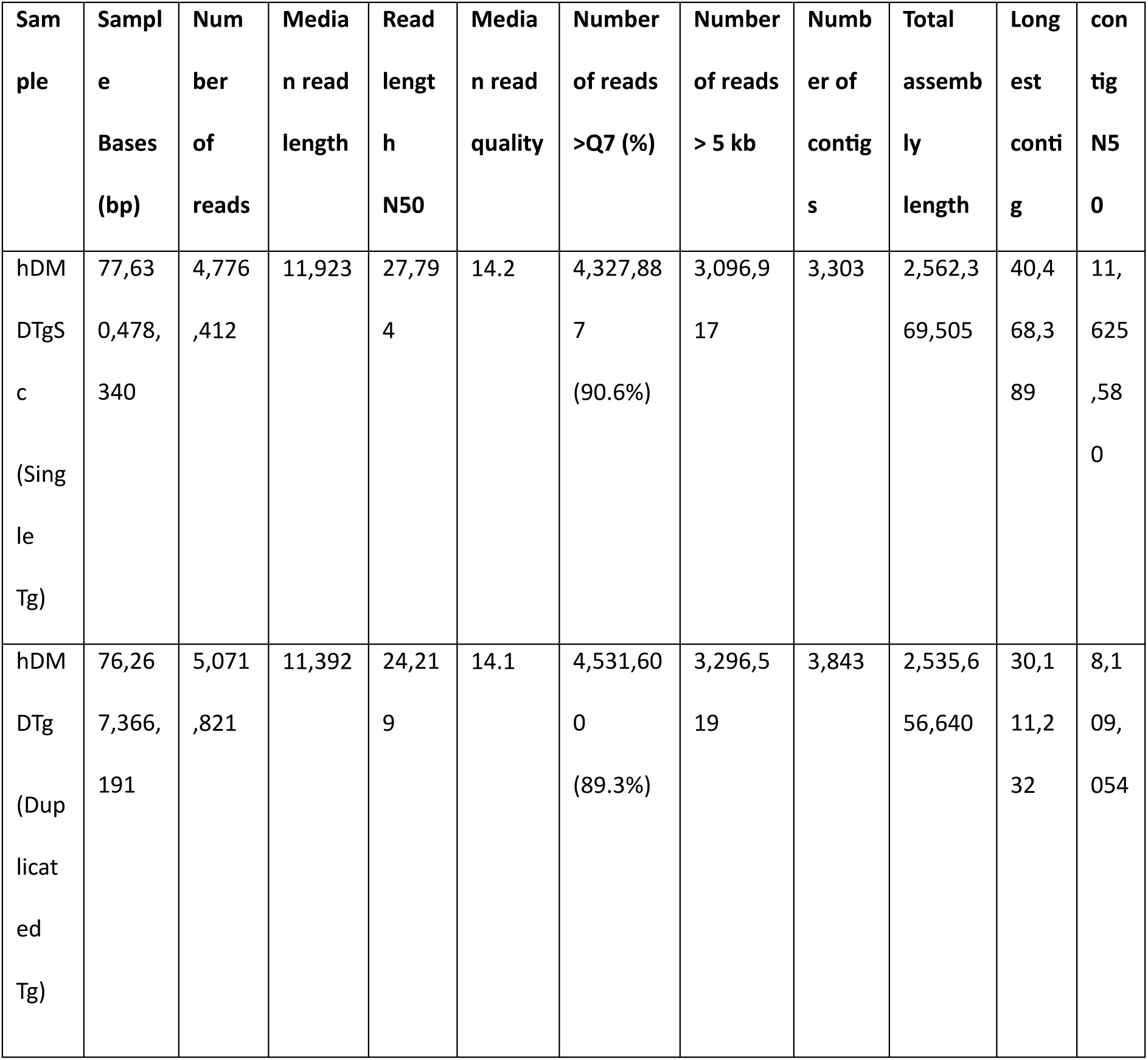
ONT output and assembly statistics.

**Supplemental Table 2:**
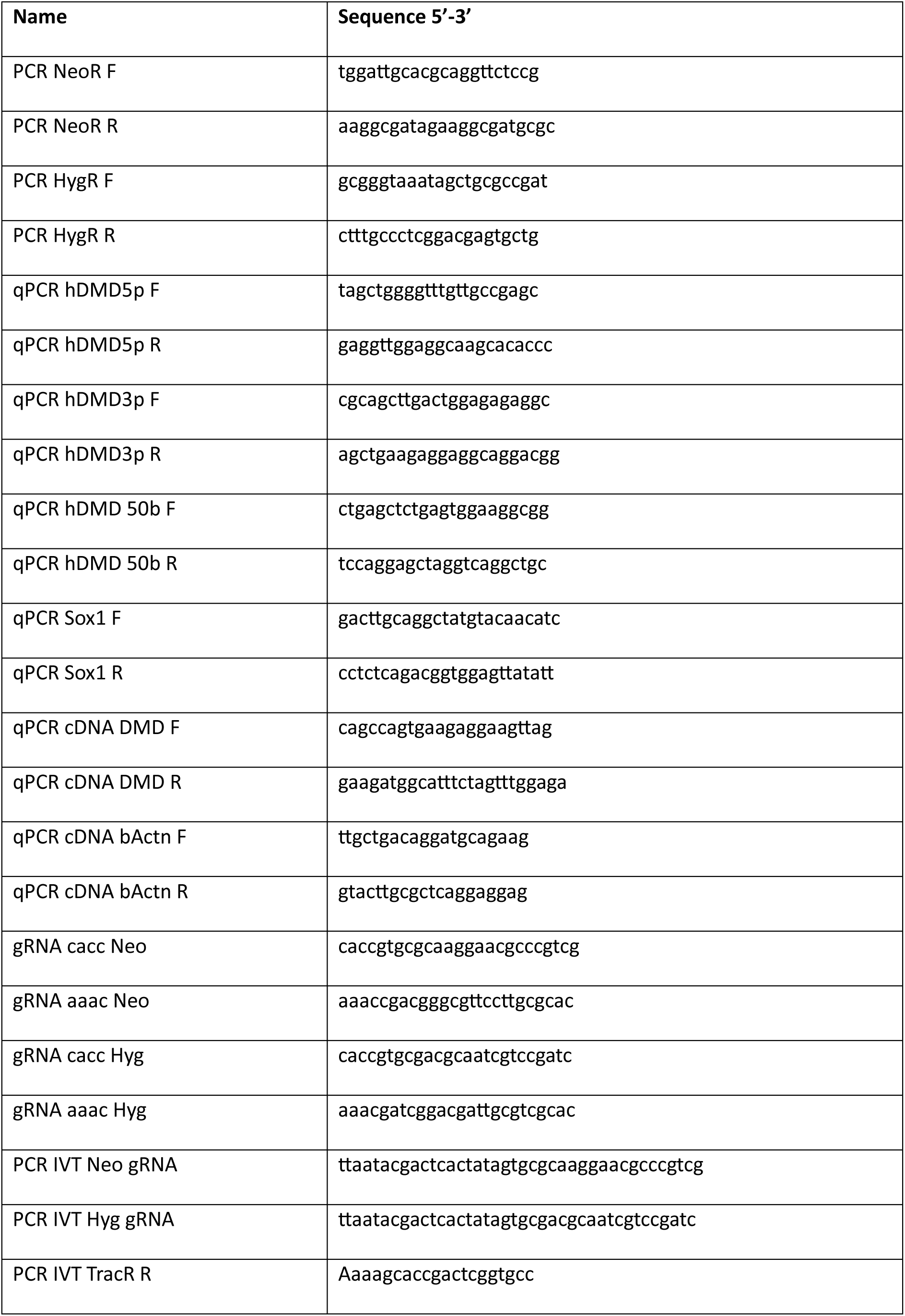
DNA primers and oligos.

**Supplementary Figure 1:**
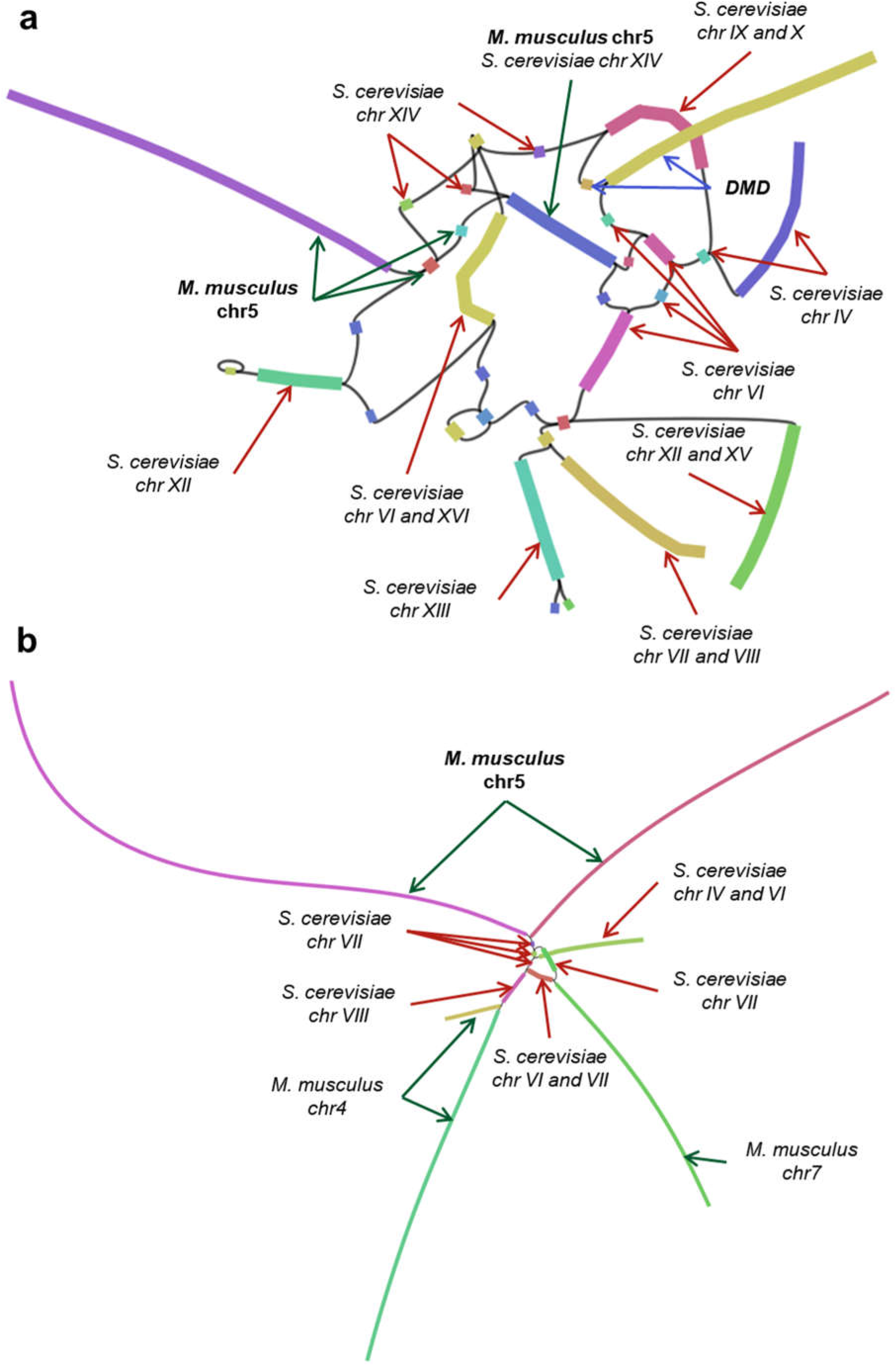
(A) Assembly sequence graph generated by Shasta of the sequences linked to the human *DMD* locus. Each of the thicker coloured nodes represents a different contig from the assembly. Labels indicate the best-matched identities of four sampled sequences of approximately 1.5 kb from the beginning, middle and end of each contig to BLAST searches of the non-redundant nucleotide database from NCBI. The branched paths in the graph indicate the different sequences at the beginning and end each copy of the DMD transgene. (B) The structure of the single copy hDMD transgene integration site. Assembly sequence graph generated by Shasta of the sequences linked to the mouse chromosome 5 insertion site (5: 133,626,547). Each of the thicker coloured nodes represents a different contig from the assembly. Labels indicate the best-matched identities of four sampled sequences of approximately 1.5 kb from the beginning, middle and end of each contig to BLAST searches of the non-redundant nucleotide database from NCBI. The presence of mouse chr4 and chr7 in this graph are likely misassembled.

**Supplementary Figure 2:**
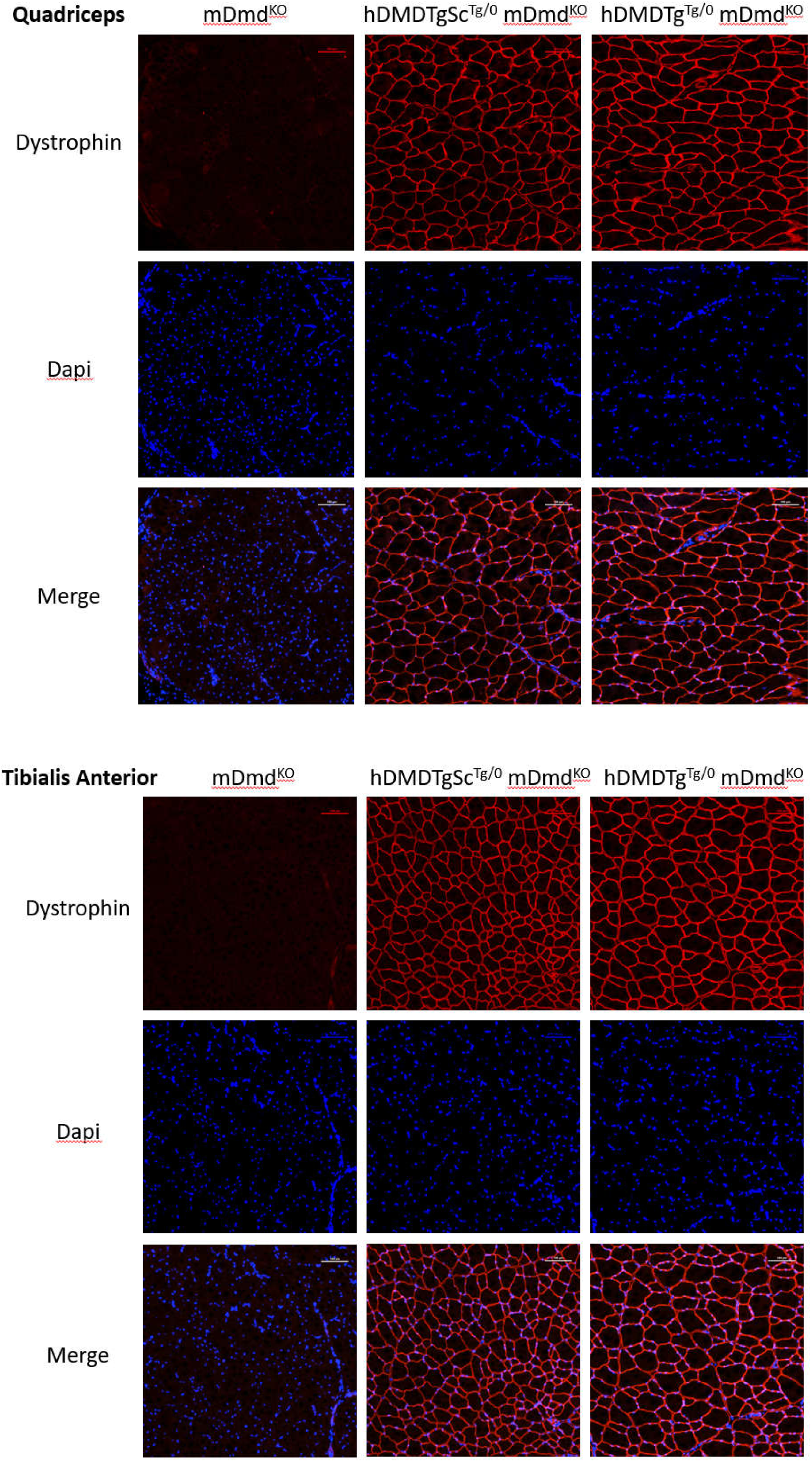
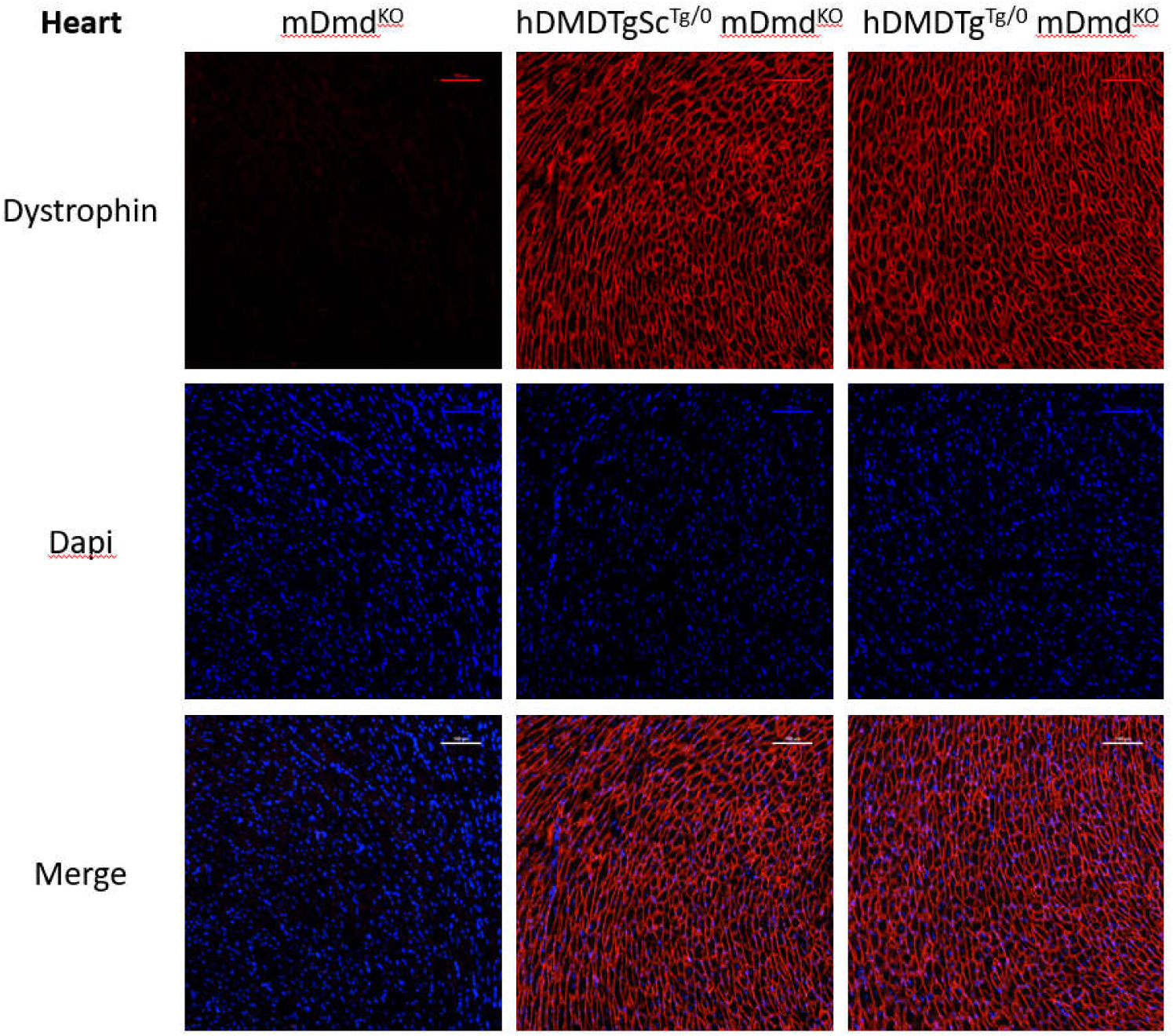
Dapi and merged immunofluoresence images. Scale bar 100 µm.

